# An advantage for targets located horizontally to the cued location

**DOI:** 10.1101/740712

**Authors:** John Clevenger, Pei-Ling Yang, Diane M. Beck

## Abstract

Over the years a number of researchers have reported enhanced performance of targets located horizontally to a cued location relative to those located vertically. However, many of these reports could stem from a known meridian asymmetry in which stimuli on the horizontal meridian show a performance advantage relative to those on the vertical meridian. Here we show a horizontal advantage for target and cue locations that reside outside the zone of asymmetry; that is, targets that appear horizontal to the cue, but above or below the horizontal meridian, are more accurate than those that appear vertical to the cue, but again either above or below the horizontal meridian (Experiments 1 and 4). This advantage does not extend to non-symmetrically located targets in the opposite hemifield (Experiment 2), nor horizontally located targets within the same hemifield (Experiment 3). These data raise the possibility that display designs in which the target and cue locations are positioned symmetrically across the vertical midline may be underestimating the cue validity effect.

## 1. Introduction

Research in a number of domains has raised the possibility that attention is preferentially allocated in a horizontal direction (Carrasco, Talgar, & Cameron, 2001; Mackeben, 1999; Montaser-Kouhsari & Carrasco, 2009; Pilz, Roggeveen, Creighton, Bennett, & Sekuler, 2012; Tse, Sheinberg, & Logothetis, 2003; Zénon, Hamed, Duhamel, & Olivier, 2009). One explanation for such findings is that these results simply reflect a much studied horizontal– vertical anisotropy (HVA) in visual performance (see Abrams, Nizam, & Carrasco, 2012 for a summary). Indeed, this visual performance explanation is consistent with increased cone density along the horizontal meridian (Curcio, Sloan, Kalina, & Hendrickson, 1990; Sawides, deCastro, & Burns, 2017) and greater numbers of neurons representing the horizontal meridian (than the vertical meridian) in the parvocellular layers of the LGN (Connolly & VanEssen, 1984) and in V1 (Tootell, Switkes, Silverman, & Hamilton, 1988; VanEssen, Newsome, & Maunsell, 1984). Another possible explanation, however, is that attention is preferentially allocated in a horizontal direction because of a bias to make horizontal shifts of attention. Such a hypothesis can be derived from the tight relationship between the oculomotor system and attention (see Smith & Schenk, 2012 for review), and in particular the fact that impairments of both midbrain and cortical structures involved in eye-movements impact exogenous orienting of attention (Gabay, Henik, & Gradstein, 2010; Rafal, Posner, Friedman, Inhoff, & Bernstein, 1988; Sereno, Briand, Amador, & Szapiel, 2006). Moreover, saccades are known to be biased in a horizontal direction in both adults (Foulsham, Kingstone, & Underwood, 2008; Gilchrist & Harvey, 2006; Tatler & Vincent, 2008) and infants (Van Renswoude, Johnson, Raijmakers, & Visser, 2016). Taking these two facts together, one might suppose that attention may be biased to move horizontally. Distinguishing between these two explanations of superior effects of attention along a horizontal axis has been hindered by the fact that the vast majority of studies confound the horizontal elongation of attention with the horizontal meridian. Here we report experiments that remove such a confound and thus support a horizontal bias in attention that does not depend on the meridian-based HVA.

### 1.1 The horizontal-vertical anisotropy

Sensitivity and acuity to stimuli that appear at some eccentricity on the vertical meridian is significantly reduced compared to sensitivity to stimuli that appear at the same eccentricity on the horizontal meridian (Abrams et al., 2012; Cameron, Tai, & Carrasco, 2002; Carrasco et al., 2001; Millodot & Lamont, 1974; Pointer & Hess, 1989; Rijsdijk, Kroon, & van der Wildt, 1980). This asymmetry falls off with distance from the horizontal and vertical meridians; half way in between the horizontal and vertical meridians (i.e. 45° polar angle), sensitivity is approximately halfway in between the sensitivity to isoeccentric stimuli on the vertical and horizontal meridians (Abrams et al., 2012). The HVA has also been observed for visual search (Carrasco, Giordano, & McElree, 2004; Chaikin, J. D., Corbin, H. H., & Volkmann, 1962; Zénon et al., 2009), letter recognition (Mackeben, 1999), short-term memory tasks (Montaser-Kouhsari & Carrasco, 2009), and change blindness task (Tse et al., 2003).

### 1.2 A horizontal bias for attention

A number of researchers have suggested that, not only might visual sensitivity be superior along the horizontal meridian, but attention may also be preferentially allocated in a horizontal direction (Mackeben, 1999; Tse et al., 2003; Zénon et al., 2009). For example, Tse, Sheinberg, and Logothetis (Tse et al., 2003) used an exogenous cueing paradigm and change detection task to map out the spatial distribution of attention. They had participants complete more than 25,000 trials over several months, allowing them to probe 149 locations within a circle around fixation that had a diameter of 25° of visual angle. On each trial participants saw a dense array of squares within the circle, followed by a blank, and then the display reappeared. On each trial, one new square appeared on the second display that was not present on the first display. Participants were asked to indicate whether the new square was red or green. Importantly, on cued trials, after the first display had been on for 506 ms, a white square (comparable in size to the target squares) appeared in 1 of 4 positions on the screen. This exogenous cue flashed twice in quick succession at 6.25° and 6.87° degrees along the cardinal axes of the display. The cues were completely uninformative in that the subsequent targets (a changed square) randomly appeared across the visual field. Nonetheless, the researchers found that change detection was improved not only at the cued location but also in an area that extended from the cued location through fixation and into the opposite visual field from the cue. This improvement at uncued locations was much more pronounced in the horizontal direction than the vertical direction, prompting researchers to propose that attention was not only allocated to the cued location but also to a symmetrical location in the other hemisphere.

Unfortunately, because the cues only appeared along the horizontal and vertical meridians, it is possible that Tse and colleagues’ (2003) result could be explained by the HVA. Indeed, Carrasco, Talgar, and Cameron (2001) showed that not only was the HVA apparent in an exogenous cueing task, attention did not interact with the asymmetry. In other words, although attention improved performance it did so equivalently on both the vertical and horizontal meridian.

There is, nonetheless, some reason to believe that there may be a horizontal bias to attention that does not rely on the HVA. These results come from the object-based cueing literature. There are now two papers (Barnas & Greenberg, 2016; Pilz et al., 2012) demonstrating a horizontal benefit to cueing, although both, interpret this horizontal advantage to the object-based component of cueing, despite the fact that the horizontal advantage persists whether or not the horizontally located target appears on the same object or not. Importantly, in these experiments, horizontalness and verticalness do not coincide with the horizontal meridians.

Both papers ran variants of the classic Egly, Driver & Rafal (1994) object-based attention task. In the classic task (Egly et al., 1994), two rectangles are displayed on either side of fixation (sometimes the rectangles are above and below fixation and sometimes they are to the left and right of fixation). Participants fixate while the rectangles are on the screen. Then, a brief exogenous cue appears at the end of the one of the two rectangles. After a delay, a target appears and participants are asked to respond. In valid trials, the target appears in the cued location. In invalid trials, the target can appear either at the other end of the cued rectangle or at the location on the other rectangle that is equidistant (and isoeccentric) from this location. The classic effect is that, while participants take longer to respond on invalid trials than on valid trials, they also take longer to respond on invalid trials where the target appears on the uncued objects as opposed to invalid trials where the target appears on the cued objects. In other words, there appears to be a benefit when a target appears on the cued object as opposed to an uncued object. This is interpreted as evidence for object-based attention, where attending to one part of object causes attention to spread to the rest of the object.

In the Pilz et al. (2012) task participants had to either identify a target (a ‘T’ or an ‘L’) appearing at the same locations as the Egly et al (1994) task. In one version of the task, 3 distractors (rotated ‘F’s) appeared, at the same time as the target, at the other 3 ends of the rectangles. While they found robust space-based attention effects, they found small and inconsistent evidence for object-based effects. More importantly to the issue investigated here, the effect varied dramatically depending on the orientation of the rectangles. When the rectangles were horizontally arranged (above and below fixation) there was a larger, more robust object-based effect (42 ms). When the rectangles were arranged vertically, they found a smaller (−-19 ms) opposite effect: a cost for invalidly cued targets that appeared on the cued object compared to the uncued object. In other words, they observed a horizontal advantage even when the target and cue appeared on different objects. The researchers found similar results in a follow-up study: same-object benefits for both accuracy and response time when the objects were arranged horizontally and same-object costs when the objects were arranged vertically.

Given that the supposed “object-based” effect reverses when the objects are arranged vertically, it seems likely that the orientation difference in object-based attention found by Pilz et al (2012) results from a general horizontal bias for attention, rather than an object-based effect per se. They consistently see an advantage for targets appearing horizontal to the cue; when that coincides with the same object (horizontally oriented objects) then the effect also looks like an object-based cueing effect. However, when the horizontal advantage is pitted against the object-based effect, as when the objects are oriented vertically, then not only do they fail to observe an object-based effect, they observe the opposite consistent with a pure horizontal advantage.

Barnas & Greenberg (2016) describe a similar horizontal advantage in their object-based task that cannot be attributed to the HVA. Their task was similar to that of Pilz et al. (2012), except that their “objects” were two grey rectangles put together to form an ‘L’. Cues appeared at the vertex of the L’s. After the vertex of the object was cued, the target could either appear at the vertex (a valid trial), at the end of the object that extended horizontally from the vertex (an invalid-horizontal trial), or at the end of the object that extended vertically from vertex (an invalid-vertical trial). They found an advantage for targets that appeared along the horizontal arm of the L compared to those that appeared along the vertical arm of the L. Because the L-shaped object was centered such that it lined up with three vertices of a square centered around fixation, these results are comparable to those of Pilz et al., (2012). In a second experiment, Barnas & Greenberg showed that this horizontal advantage persisted even when the cue and target appeared on different objects. In other words, both Pilz et al. 2012 and Barnas & Greenberg, 2016 show an advantage for targets appearing horizontally (as opposed to vertically) to a cue regardless of whether the cue and target appear on the same object or different objects. Both results were interpreted as evidence that attention can move more easily along a horizontally oriented object than a vertical one, but this would not explain why attention seems to move more easily in a horizontal direction even when that means jumping from one object to another (Pilz et al., 2016). Instead, we suggest that both studies have in fact provided evidence for a horizontal advantage that is independent of whether attention moves along an object or not^1^. Thus, here we ask whether this effect may not be restricted to object-based attention but instead may reflect a general bias for attention to shift horizontally, by removing the additional objects.

### 1.3 Horizontal eye-movements and attention

There is another reason to believe that attention could have a horizontal bias. A number of researchers, starting most notably with Rizzolatti, Riggio, Dascola, and Umilta (1987), have argued that attention and eye-movements are linked. Preparing to make an overt eye-movement involves many of the same brain regions as covertly moving attention (Smith & Schenk, 2012). Moreover, saccades are not only preceded by shifts of attention (Deubel & Schneider, 1996; Hoffman & Subramaniam, 1995; Kowler, Anderson, Dosher, & Blaser, 1995; Shepherd, Findlay, & Hockey, 1986), but patients who have difficulty executing saccades show concomitant deficit in covertly orienting to exogenous cues (Gabay et al., 2010; Rafal et al., 1988; Sereno et al., 2006). It is thus reasonable to assume that biases present in the eye-movement system may be replicated in the attention system.

Horizontal eye-movements are more common than other directions in both adults (Foulsham et al., 2008; Gilchrist & Harvey, 2006; Tatler & Vincent, 2008) and infants (Van Renswoude et al., 2016). The genesis of this bias may be biomechanical; horizontal eye-movements only require one pair of muscle movements whereas oblique or vertical saccades require multiple pairs (Becker & Jürgens, 1990; Henn & Cohen, 1973; Viviani, Berthoz, & Tracey, 1977). Another possibility is that the same physiological factors that lead to the HVA, namely the overrepresentation of the horizontal as opposed to vertical dimension, in the eye (Curcio et al., 1990; Sawides et al., 2017) and brain (Connolly & Van Essen, 1984; Tootell et al., 1988; Van Essen et al., 1984), could produce a bias to sample in the horizontal direction. A third possibility is that the distribution of objects in the world tends to occur along the horizon and we have therefore adapted our eye-movements accordingly. Finally, the bias may simply reflect the fact that our visual field has a greater extent in the horizontal direction than it does in the vertical, by virtue of having two horizontally displaced eyes, making information presented along the horizontal dimension more likely to induce an eye-movement. Regardless of the cause of the horizontal eye-movement bias, it is possible that attentional shifts have inherited this bias.

Interestingly, to our knowledge, the only experiments that have demonstrated this horizontal advantage have employed an exogenous attention task. Rizzolatti et al., (1987) employed an endogenous attention task and reported a numerical advantage for vertically located targets, although this difference did not reach significance. The fact that the horizontal advantage has been observed only with exogenous attention tasks is again suggestive of an oculomotor influence, as patients with oculomotor deficits have more consistently shown deficits in exogenous attention than in endogenous attention (Gabay et al., 2010; Rafal et al., 1988; Sereno et al., 2006; Smith, Rordan, Jackson, 2004—although see Craighero, Carta, Fadiga, 2001 for exception).

### 1.4 Current Study

Here we ask whether such a horizontal bias exists for attention in the absence of eye-movements, while controlling for the HVA. Specifically, we ask whether exogenous cues induce a greater bias to detect a target in a position located along a horizontal trajectory from the cue than an equivalently distant position located along a vertical trajectory from the cue. Critically, all cues and targets are located outside the zone of the HVA (i.e. maximally distant from both the vertical and horizontal meridian; at 45° polar angle).

## 2. Experiment 1

In Experiment 1, we had participants complete a visual search task in which they were instructed to covertly find a face among non-faces. The target array (consisting of 1 target face and 2 non-target non-faces) was preceded by a spatial precue. The cue onset before the target array and its validity as a spatial cue was manipulated. Targets and cues could appear, with equal probability, in one of four positions arranged in a square around fixation, such that all four positions fall outside of the zone of the Horizontal-Vertical Asymmetry (Figure 1). On 50% of trials, the target appeared exactly where the cue had appeared (a valid trial) and on 50% of trials the target appeared in an uncued location positioned either vertically or horizontally from the cues. Cues could appear at any of the four locations on the square and, when the cue was invalid, the target could either appear horizontal to the cue across the vertical midline (invalid-horizontal) or vertical to the cue across the horizontal midline (invalid-vertical). Targets appeared equally often at all locations in all conditions, and thus no location biases were introduced by the task parameters that should cause participants to preferentially allocate their attention to one location over the other (except for the presence of the cue). Critically, there were no targets on the horizontal midline, and the distance between the targets and cues in the invalid-horizontal position was equivalent to the distance between targets and cues in the invalid-vertical condition. In order to test whether there is a performance advantage for targets appearing horizontal to the cue we subdivided the invalid trials into two basic kinds: those where the target subsequently appeared horizontal to the cue and those where the target appeared diagonal to the cue.

**Figure 1:**
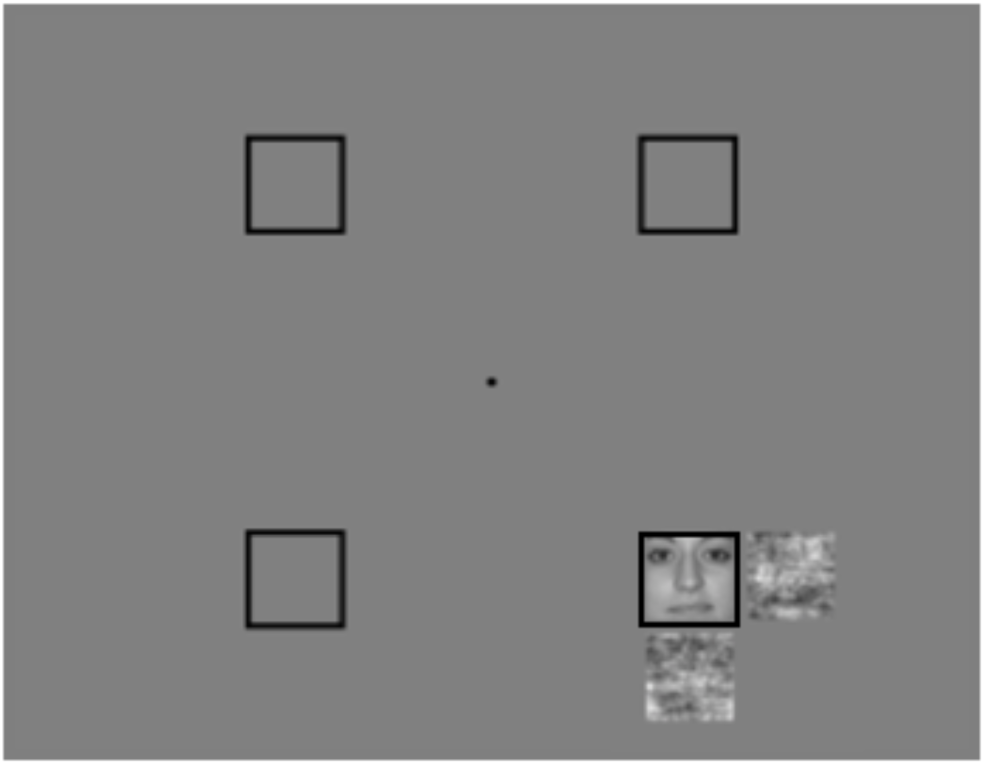
Possible cue and target locations from Experiment 1, with example target and flankers in one position. Both the cue and target appeared equally often at all 4 positions, although once a cue location was chosen the target could only appear either at that location (50% of trials) or vertically (25% of trials) or horizontally (25% of trials) displaced from that position (i.e. could not appear diagonally from target).

### 2.1 Material and Methods

#### 2.1.1 Participants

Twelve participants were recruited from the University of Illinois at Urbana-Champaign in exchange for credit in an introductory Psychology course. All 12 gave informed consent based on the University of Illinois IRB protocol and reported normal or corrected-to-normal vision and color vision.

#### 2.1.2 Apparatus, Stimuli, and Procedure

Participants were seated 57 cm (chinrest enforced) from a 24-inch LCD monitor set to a refresh rate of 100 Hz. All stimuli were derived from a single image taken from a publicly available database (Minear & Park, 2004). This image, used as the target, was a square cutout of a grayscale, Caucasian female face subtending 1.5° × 1.5° of visual angle. Masks and flankers were 100% phase-scrambled versions of this image. The participants’ task was to determine whether the central face was right side up or inverted. Target locations were cued, with 50% validity, by the appearance of a box.

The cue and target locations were positioned on an imaginary square centered at fixation (Figure 1). The vertices of the square were located 5° of visual angle from fixation. The distance between any two vertices along a side of the imaginary square was about 7.1° of visual angle. Starting from the upper half of the vertical midline (which would be 0° of polar angle) the vertices appeared at 45°, 135°, 225°, and 315° of polar angle, respectively.

The trial sequence is depicted in Figure 2. As each trial began, participants saw a black dot at fixation which remained on a middle gray screen throughout the trial. Then, 300 - 500 ms after the onset of the fixation cross, a spatial precue appeared for 50 ms before offsetting. The precue was a black outline square subtending 2.25° × 2.25° of visual angle. After the precue, the screen was blank for 50 ms before the onset of the target array. The target array, consisting of the target and two non-targets, appeared for 100 ms (Figure 2). The target was always the same square face and the non-targets were different phase-scrambled versions of this face. The two phase-scrambled non-targets flanked the target such that they always appeared on the two sides of the target facing away from fixation (see Figure 2). For instance, if the target appeared at the top right vertex of the square, the non-targets would appear above and to the right of the target. If the target appeared at the bottom left of the vertex, the non-targets would appear below and to the left of the target.

**Figure 2:**
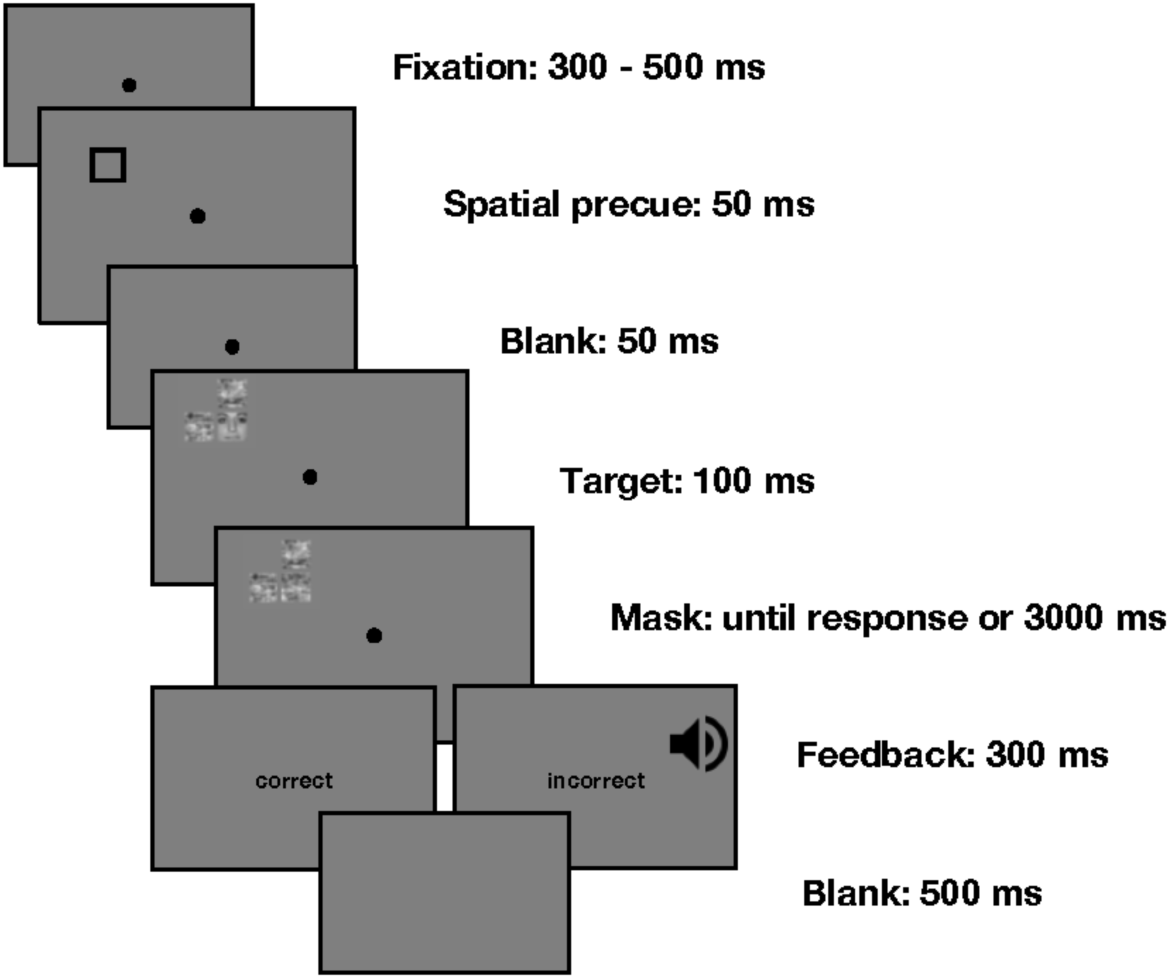
Trial sequence for Experiment 1. Cue and target locations were drawn equally often from the locations presented in Figure 1.

In a pilot study, we discovered that these flankers are actually necessary for obtaining a sizable validity effect in accuracy, with the potential detect a modulation of the validity effect by location (horizontal vs vertical). We believe this is because it is necessary that the flankers interfere/compete (Beck & Kastner, 2009; Desimone & Duncan, 1995; Kastner et al., 2001; Kastner, DeWeerd, Desimone, & Ungerleider, 1998) with the target in order for the target to be unresolvable and thus result in errors, as opposed to just slower reaction time. This competition, coupled with the short exposure time before masking (100 ms), should reduce the amount of information the visual system can quickly extract about the target and help make differences in attentional deployment (used to resolve the target) more obvious. We note that we refer to these non-targets as flankers simply because they appear on both sides of the target; they did not have a response associated with them and so they were not flankers in the tradition Eriksen flanker paradigm sense (Eriksen & Eriksen, 1974).

Cues were valid on 50% of trials and invalid on 50% of trials. When invalid, the target could appear in one of two positions: either on the vertex directly horizontal to the cue or on the vertex directly vertical to the cue. When invalid, the target never appeared at the vertex diagonal to the cue.

The target appeared right-side up on half of the trials and upside down on the other half. After 100 ms, three masks appeared in the same locations as the items on the target array. These masks were phase-scrambled versions of the target face and were always different from both the non-targets and each other. The masks stayed on the screen until either a response was made or 3000 ms had passed since the target array appeared. If the participant got the trial correct, the word ‘correct’ would appear at fixation for 300 ms, replacing the fixation cross. If the participant got the trial incorrect, the word ‘incorrect’ would appear at fixation along with a loud tone for 300 ms. Next, a 500 ms black screen appeared before the next trial started with a fixation cross.

Participants were instructed to covertly search the target array and indicate via keypress whether the target face on a given trial was upright (the ‘z’ key) or upside down (the ‘?/’ key). Participants were told to respond quickly but without sacrificing accuracy and were strongly encouraged to keep their eyes at fixation during trials. Trials without responses were terminated after 3000 ms and marked incorrect. Participants completed 1 practice block of 64 trials (not included in analysis) and 6 experimental blocks of 64 trials each. After each block, participants took a self-timed break. During the break their accuracy on the just-completed block was displayed along with their highest accuracy score on any previously completed block and a reminder to keep their eyes at fixation during the trials.

The predictions for Experiment 1 are straightforward. First, we expect to see a performance advantage for the valid location relative to either invalid location, as has been seen many times before. More importantly, we are interested in performance at the invalidly cued locations. Previous research indicates that there is an attentional benefit for targets appearing horizontal to an invalid cue (Carrasco, Talgar, & Cameron, 2001; Mackeben, 1999; Montaser-Kouhsari & Carrasco, 2009; Pilz, Roggeveen, Creighton, Bennett, & Sekuler, 2012; Tse, Sheinberg, & Logothetis, 2003; Zénon, Hamed, Duhamel, & Olivier, 2009). If this benefit is due to increased sensitivity along the horizontal meridian, in this experiment, where no target or cue appears on the meridian, we should see equivalent performance for the horizontally located invalid position as the vertically located invalid position. If, however, the horizontal advantage is more general we should see improved performance for targets located horizontally from the cue (invalid-horizontal) relative those located vertically (invalid-vertical). The task was designed to produce accuracy differences as a function of the cue and location, but we will also explore RT effects.

### 2.2 Results

#### 2.2.1 Inferential statistics

Accuracy and response time data are presented in Table 1 and in Figure 3. The valid condition produced more accurate (Mean difference = 9.7%, SD = 7.5%, 95% CI of the mean = [4.9%, 14.4%], dz = 1.29, 95% CI of the effect size = [0.47, 2.12]) and faster responses (Mean difference = −98.4 ms, SD = 85.2 ms, 95% CI of the mean = [−152.96, −43.89], dz = −1.15, 95% CI of the effect size = [−1.62, −0.75]) than the invalid condition. To test the hypothesis that there is an attentional benefit for targets horizontal to an invalid cue we computed a difference score for each participant by subtracting the participant mean in the invalid-vertical condition from the participant mean in the invalid-horizontal condition (Figure 3B). This difference score represents the gain in accuracy for a given participant when the target appeared horizontally to the invalid cue compared to when it appeared vertically. All reported confidence intervals are bias-corrected, accelerated bootstrapped confidence intervals (Efron & Tibshirani, 1993) calculated using the wBoot R package (Weiss, 2016). The difference score will be positive when a participant’s accuracy is higher in the invalid-horizontal condition than in the invalid-vertical condition. The group difference scores show a large benefit for horizontal, relative to vertical, targets after invalid cues (M = 9.8%, SD = 8.2%, 95% CI of the mean = [5.8%, 14.7%], dz = 1.2, 95% CI of the effect size = [0.71, 1.66]).

**Figure 3:**
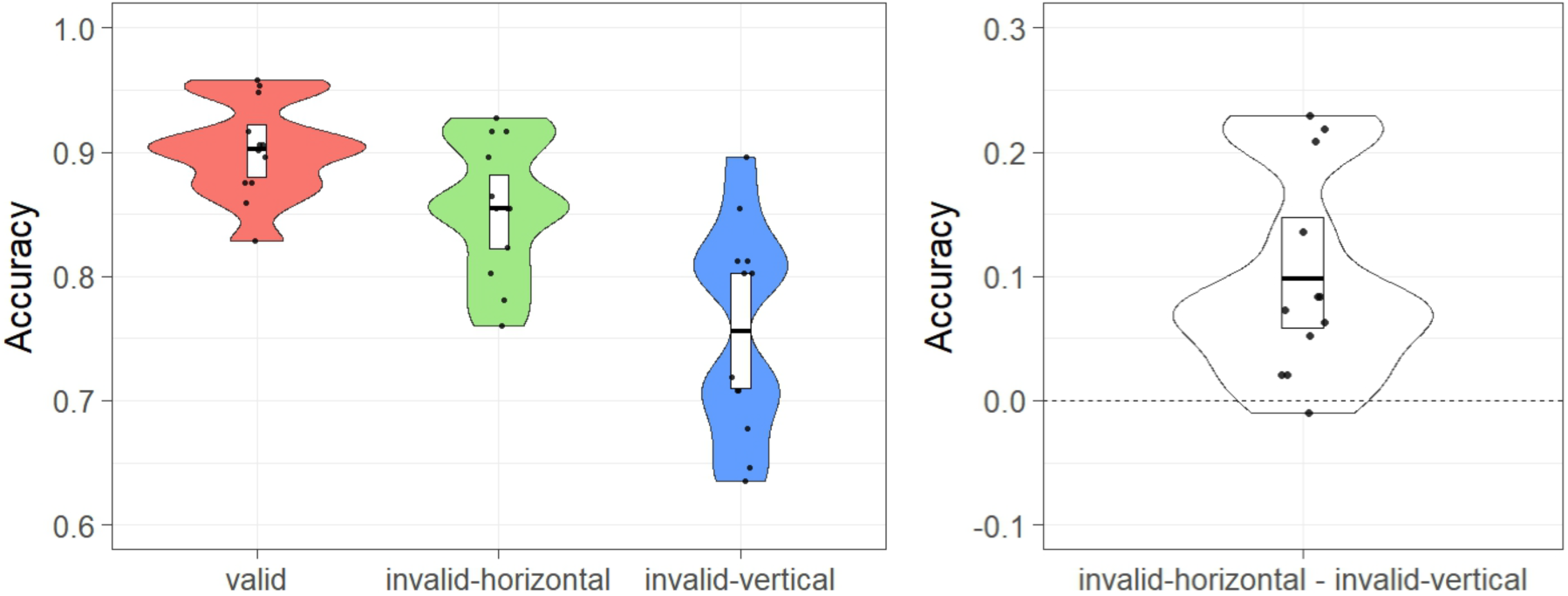
Accuracy data from Experiment 1. Each dot represents one participant, and is slightly jittered to the left or right for readability. The distributions are kernel density plots of participant-level difference scores. The white bars represent BCa bootstrapped 95% confidence intervals (Efron & Tibshirani, 1993). The horizontal line in the center of each white bar is the group mean. Each dot represents one participant, and is slightly jittered to the left or right for readability. The distributions are kernel density plots. A) accuracy as a function of condition. B) Participant-level accuracy differences between the invalid-horizontal and the invalid-vertical conditions. The y-axis represents how much higher accuracy was in the invalid-horizontal condition compared to the invalid-vertical condition.

**Table 1.**
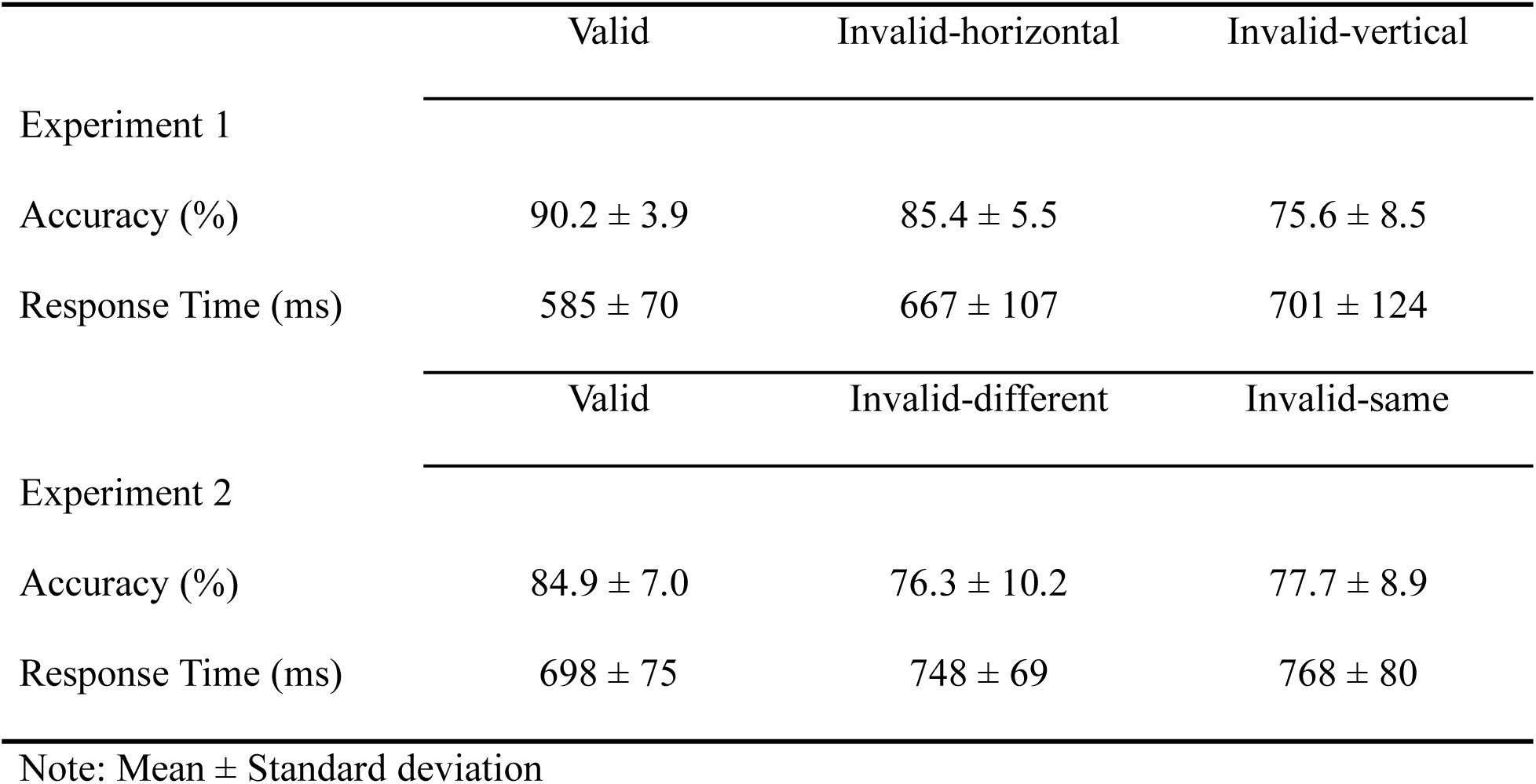
Descriptive statistics for Experiments 1 and 2

A Monte Carlo randomization test (permutation test) was conducted to determine whether this observed benefit was greater than chance. Assuming the null is true (that the invalid-horizontal and invalid-diagonal conditions come from the same distribution), then for each participant their observed score in one of the conditions is equally likely to have been observed in the other condition. To simulate this, we conducted 100,000 iterations where on every iteration each participant had a 50% chance that the condition labels on their two scores were switched. After going through each participant and either switching or not switching their condition labels, differences scores were again computed and the mean difference score was taken. This process was repeated 100,000 times and 100,000 difference scores were collected. If there is no actual difference between the two conditions, then the difference score we observed in our data (9.8%) should not be extreme compared to the distribution of difference scores generated in the randomization test. Of the 100,000 iterations run, 42 produced a difference score with an absolute value as great or greater than observed, 9.8%, thus resulting in p < .001 (Phipson & Smyth, 2010). The same procedure was performed on the difference between valid and invalid conditions (collapsed across horizontal and vertical conditions) resulting in 142 difference scores with an absolute value as great or greater than observed (i.e. 9.7%), corresponding to p < .01.

#### 2.2.2 Exploratory RT analyses

While the primary analysis concerns accuracy, difference scores were computed for the response times of the two conditions of interest. We subtracted each participant’s response time in the invalid-horizontal condition from their response time in the invalid-vertical condition. The differences scores revealed that responses in the invalid-vertical condition took perhaps a little longer than those in the invalid-horizontal condition (M = 34 ms, SD = 45 ms, 95% CI of the mean = [13 ms, 64 ms], dz = 0.75, 95% CI of the effect size = [0.32, 1.23]). A randomization test (using the same procedure as above) was conducted. Of the 100,000 iterations, 2282 produced a difference score with an absolute value as great or greater than the observed one (34 ms), p = 0.023. Thus, although less robust, we see a similar effect in RT as in accuracy.

### 2.3 Discussion

Following an invalid cue, participants performed significantly better when targets appeared at a location horizontal to the cue than when targets appeared in an equivalent vertical distance from the cue. Importantly, neither cue nor target appeared along the horizontal or vertical meridians, and thus this horizontal advantage cannot be attributed to the Horizontal-Vertical Anisotropy (Abrams et al., 2012). Instead, the spatial arrangement of our stimuli were more similar to that of Pilz et al (2012) and Barnas et al (2016) who showed a horizontal advantage in the context of an object-based attention paradigm; that is, object-based effects were larger when the target appeared on an horizontally oriented object, either above or below fixation, than a vertically oriented object. They, reasonably, concluded that the advantage was likely to be a property of object-based attention. However, our results suggest instead that their horizontal object-based advantage might be better explained by a general advantage for targets located horizontally to an invalid cue, regardless of whether they appear on the same object or not.

Although our display arrangement in Experiment 1 allowed us to rule out horizontal meridian effects, it introduces a new concern. The display in Experiment 1 was an imaginary square centered at fixation (Figure 1). Thus, when the cue was invalid, targets in the invalid-horizontal condition always appeared in a different visual hemifield (left or right) than the cue had previously appeared in while targets in the invalid-vertical condition always appeared in the same visual hemifield as the cue had previously appeared in. Some researchers have argued that the left and right cortical hemispheres have different pools of attentional resources (Alvarez & Cavanagh, 2005; Franconeri, Alvarez, & Enns, 2007) and that processing of difficult visual tasks is thus easier when the stimuli fall into different visual hemifields as opposed to the same hemifield (though see Clevenger &Beck, 2014 and Scalf &Beck, 2010 for alternative explanation). If true, this might mean that cues and targets in the invalid-horizontal condition of Experiment 1 were processed by separate pools or resources while the cues and targets in the invalid-vertical condition were processed by the same pool of resources. This might cause a benefit in performance for the invalid-horizontal condition relative to the invalid-vertical condition by virtue of the invalid-vertical condition being more resource intensive.

## 3. Experiment 2

In Experiment 2, we attempted to address the concern with Experiment 1 that asymmetrical hemifield load might explain the observed advantage for horizontal relative to vertical targets. We did this by altering the stimuli placement relative to Experiment 1. In Experiment 1, the stimuli appeared at the vertices of an imaginary square centered at fixation (Figure 1). This led to two invalid conditions: one where the targets appeared directly horizontal to the cue and one where the targets appeared directly vertical to the cue. In Experiment 2, we rotated the imaginary square either left or right so that both invalid conditions involved targets appearing diagonally from the cue, meaning there were no horizontal targets (Figure 4). However, we kept the asymmetrical hemifield load from Experiment 1. When an invalid cue appeared, the target could either appear (diagonally) in the same hemifield as the cue or (diagonally) in the opposite hemifield as the cue. If the advantage for horizontal targets that we found in Experiment 1 occurred because of an asymmetrical hemifield load we should find a similar difference in Experiment 1. However, if the advantage was due to an attentional benefit for horizontally presented targets following invalid cues, we should not find such a difference here.

**Figure 4:**
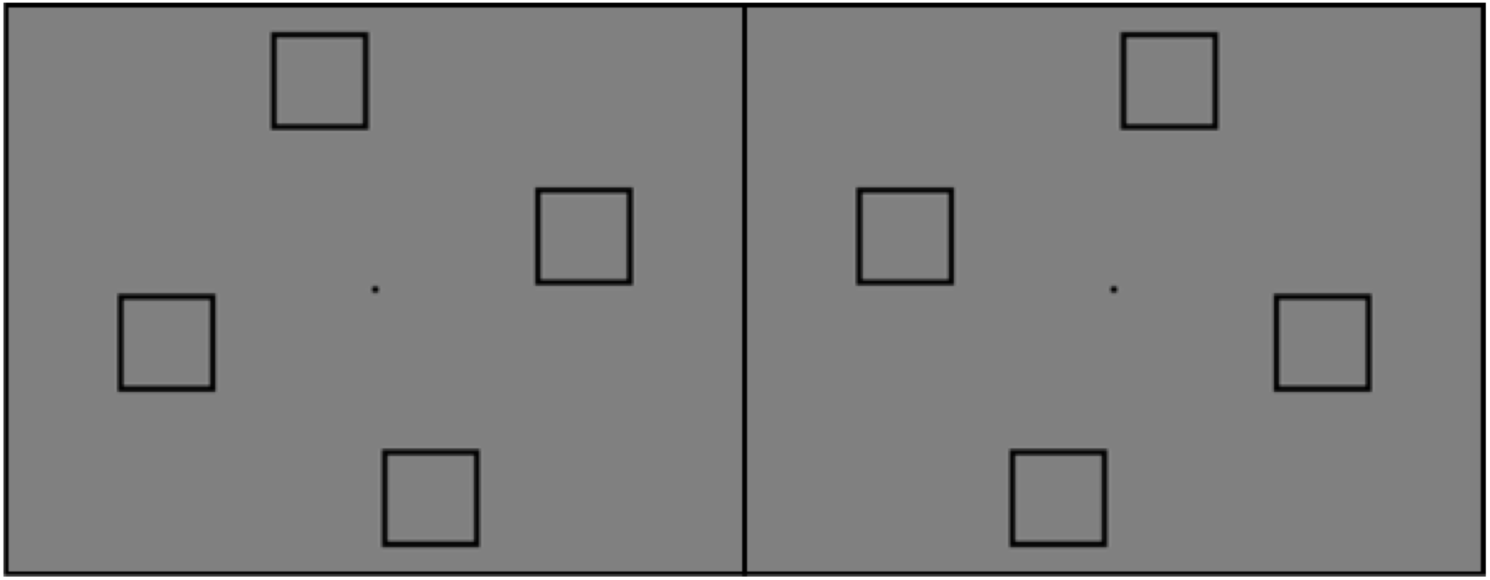
Possible cue and target locations in both types of displays used in Experiment 2. Participants were assigned to one of the two display types and all of their trials used their assigned display type.

### 3.1 Material Methods

#### 3.1.1 Participants

A post-hoc power analysis of Experiment 1 indicated that we had 97% power to detect an effect of the observed size with 12 subjects. Thus, for Experiment 2, we again set our desired sample size at 12. Thirteen participants were recruited from the University of Illinois at Urbana-Champaign in exchange for credit in an introductory Psychology course. All 13 gave informed consent based on the University of Illinois IRB protocol and reported normal or corrected-to-normal vision and color vision. One participant was excluded because they got fewer than 60% of the experimental (non-practice) trials correct. The analysis was conducted on the remaining 12 participants.

#### 3.1.2 Apparatus, Stimuli, and Procedure

Experiment 2 was identical to Experiment 1 except for the following. In Experiment 2, the display was a rotated imaginary square (Figure 4). There were two versions of the display. One version had the vertices placed (with 0° of polar angle being on the upper half of the vertical midline) at 75°, 165°, 255°, and 345° of polar angle, respectively (left panel of Figure 4). The other version had vertices placed at 15°, 105°, 195°, and 285° of polar angle, respectively (right panel of Figure 4). Each participant was assigned to one of the two display types with assignment alternating back and forth as participants came in. Participants only saw trials of one display type. The vertices of the square were again located 5° of visual angle from fixation and were the locations where the cues and targets appeared. The distance between any two vertices along a side of the imaginary square was about 7.1° of visual angle. Cues were again valid on 50% of trials and invalid on 50% of trials. When invalid, the target could appear in one of two positions: either on the vertex in the same visual hemifield as the cue or on the vertex in the opposite visual hemifield as the cue. When invalid, the target never appeared at the vertex diagonal to the cue.

#### 3.2.1 Inferential statistics

Accuracy in the display where the first vertex on the right was placed 75° from the upper vertical midline (left panel of Figure 4) (M = 81.4%, SD = 10.9%) did not substantially differ from accuracy in the other display (right panel of Figure 4) (M = 80.6%, SD = 4.4%), thus display type was collapse across subsequent analyses. The resulting accuracy and response time data are presented in Table 1 and in Figure 5. The valid condition produced more accurate (Mean difference = 7.9%, SD = 6.7%, 95% CI of the mean = [3.7%, 12.1%], dz = 1.18, 95% CI of the effect size = [0.81, 1.62]) and faster responses (Mean difference = −60.1 ms, SD = 53.2 ms, 95% CI of the mean = [−93.97, −26.29], dz = −1.13, 95% CI of the effect size = [−1.57, −0.73]) than the invalid condition.

**Figure 5:**
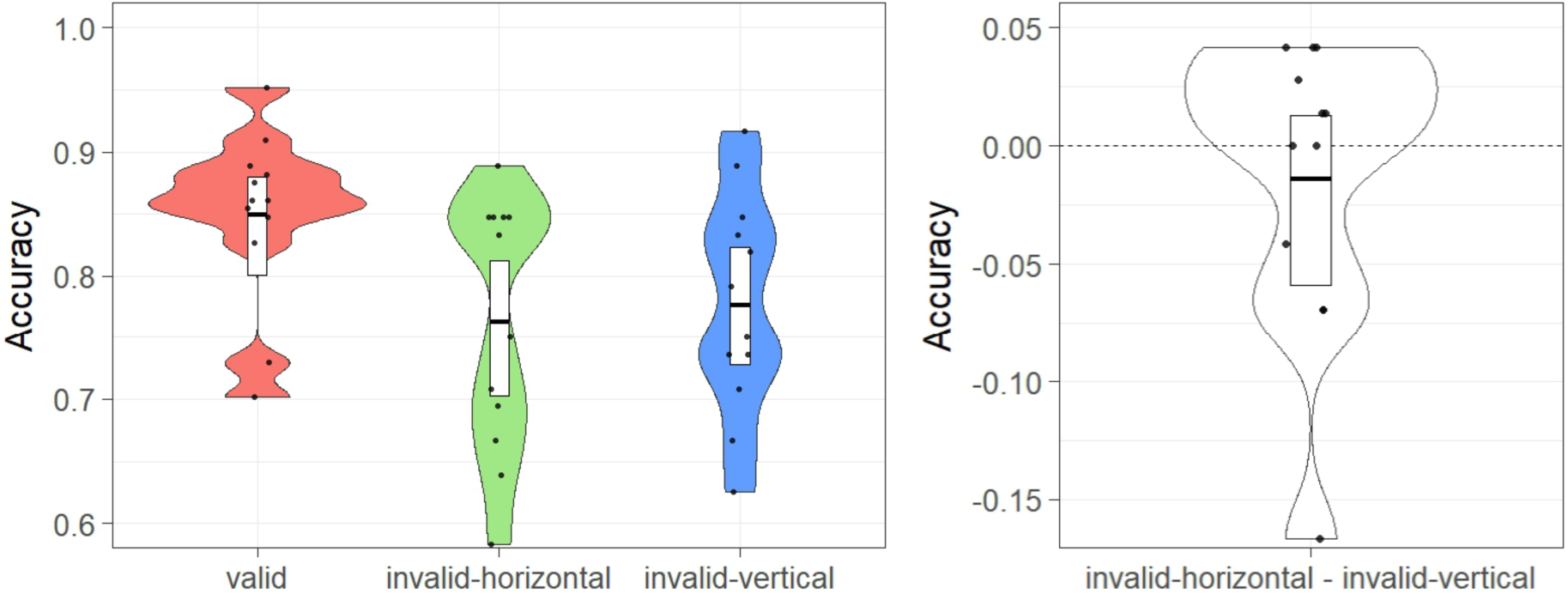
Accuracy from Experiment 2. Violin plot representation as in Figure 3. A) Accuracy as a function of condition. B) Participant-level accuracy differences between the invalid-different and the invalid-same conditions.

We again computed a difference score for each participant. We subtracted the participant mean in the invalid-same hemifield condition from the participant mean in the invalid-different hemifield condition (Figure 5B). This difference score will be positive to the extent that a participant was more accurate in the invalid-different hemifield condition than in the invalid-same hemifield condition. The group difference scores show only a minimal difference between the two conditions (M = −1.4%, SD = 6.3%, 95% CI of the mean = [−5.9%, 1.3%], dz = 0.42, bootstrapped 95% CI of the effect size = [−0.6, 0.79]). We again conducted a randomization test to determine whether the observed difference was greater than that expected due to chance. The procedure was the same as in previous experiments. Of the 100,000 iterations run, 47546 produced a difference with an absolute value as great or greater than the observed difference (−1.4%), returning p = 0.475, indicating no significant difference in accuracy as a function of whether the target was in the same hemifield as the cue or not. The same procedure was performed on the difference between valid and invalid conditions (collapsed across horizontal and vertical conditions) resulting in 0 difference scores with an absolute value as great or greater than observed (i.e. 7.9%), corresponding to p < .001, indicating a significant validity effect.

#### 3.2.2 Exploratory RT analyses

We analyzed the response time scores for both the invalid-same hemifield condition and the invalid-different hemifield condition. We again computed difference scores. We subtracted each participant’s response time in the invalid-same hemifield condition from their response time in the invalid-different hemifield condition. The differences scores revealed a small benefit for the invalid-different hemifield condition (M = 20 ms, SD = 31 ms, 95% CI of the mean = [1 ms, 35 ms], dz = 0.64, 95% CI of the effect size = [−0.21, 1.42]), which falls just outside of the confidence interval. A randomization test (using the same procedure as above and uncorrected for multiple corrections/exploratory analyses) also did not reach significance. Of the 100,000 iterations, 4858 produced a difference score with an absolute value as great or greater than the observed one (20 ms), p = 0.049.

### 3.3 Discussion

In Experiment 2 we were interested in determining whether the benefit for horizontal targets found in Experiment 1 was due to asymmetric hemifield load rather than an actual advantage for targets presented horizontal to an invalid cue. We tried to test between these two accounts by preserving asymmetric hemifield load while removing horizontal targets. When cues were invalid, some targets appeared diagonally in the same hemifield as the cue and some appeared diagonally in the opposite hemifield as the cue. If hemifield load was responsible for the effect in Experiment 1 we should have found an advantage for those targets appearing in a different hemifield for the cue, which we did not. In fact, performance when targets appeared in a different hemifield (76.3%) was effectively identical to performance when targets appeared in the same hemifield as the cue (77.7%). This increases our confidence that asymmetric hemifield load does not explain the effect observed in Experiment 1.

Experiment 2 has another property of interest: in the invalid condition, the target appeared at two levels of steepness (relative to horizontal) from the cue. If the horizontal midline is 0° of polar angle, the target appears either 60° of polar angle above or below the cue in the same hemifield condition and 30° of polar angle above or below the cue in the different hemifield condition. This might be important if there is a continuous detriment to performance the less horizontal (i.e., steeper) the shift of attention is between the cue and the target. This might be true under a variety of models of the effect. If horizontally presented targets benefit because shifting attention horizontally is easier/faster than shifting attention non-horizontally, the more horizontal component there is in a shift vector the better performance might be. Thus, we might expect performance to be better when shifting 30° than shifting 60° because there is a larger horizontal component to the 30° shift vector. However, we did not observe any accuracy difference between these differently steep shifts.

## 4. Experiment 3

Together Experiments 1 and 2 suggest there is an advantage for targets located horizontally to a cue that cannot be explained by superior sensitivity along horizontal meridian or differences in hemifield load. In Experiment 3 we ask whether locating the target in the horizontally opposite hemifield is necessary, or more simply, whether a horizontal advantage is observed within a hemifield.

Figure 6 shows the displays used in Experiment 3. The task was similar to the previous experiment. The cues could appear in any of the eight locations on the screen. The target and cue sizes were scaled to control for eccentricity and cortical magnification. Once the cue appeared, the target could either appear in the same location as the cue (a valid trial) or in one of two locations in the same hemifield as the cue: either directly horizontal or directly vertical to the cue.

**Figure 6:**
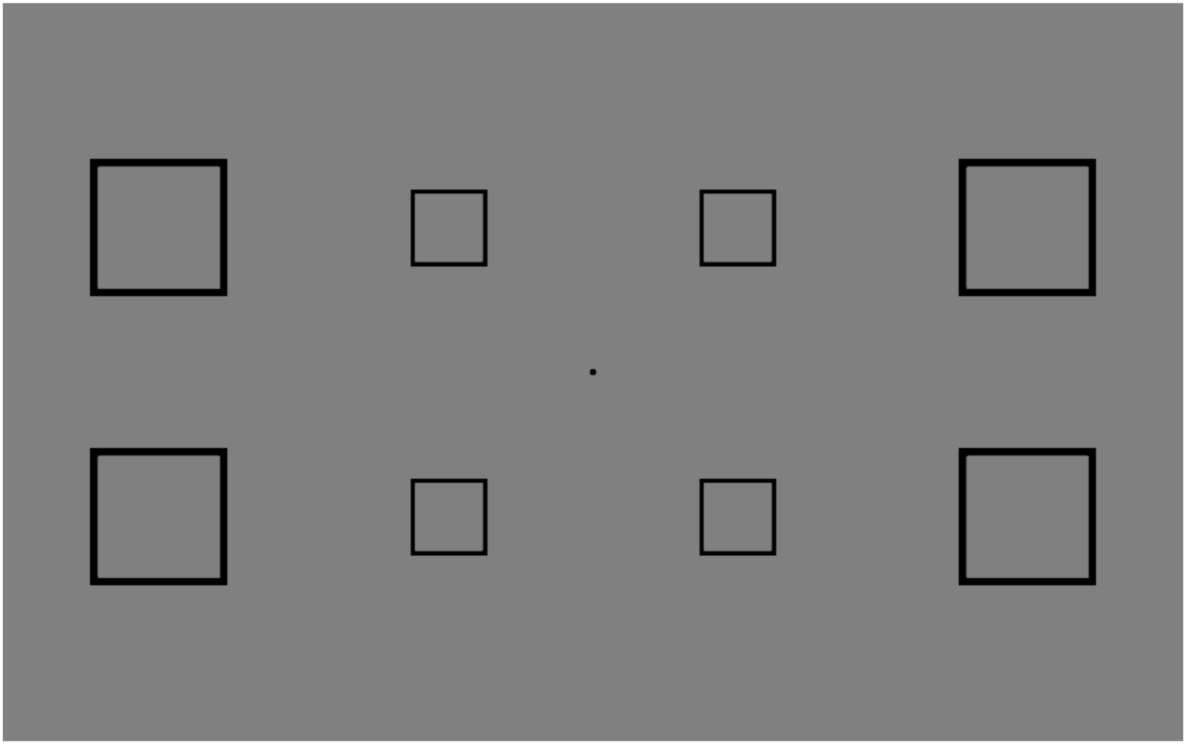
Target and cue locations in Experiment 3. Cues appeared equally often at all eight locations. When the cue was invalid, targets appeared within the same hemifield as the cue at either the location directly horizontal to or directly vertical to the cue. Outside positions are placed about 8.1° and the inside positions 4° of eccentricity relative to fixation.

### 4.1 Material and Methods

#### 4.1.1 Participants

To accommodate the possibility that the within-hemifield manipulation might result in a smaller horizontal advantage, we powered our experiment to have 80% power to detect an effect 2/3 the size of that found in Experiment 1, resulting in a desired sample size of 15. Fifteen participants were recruited from the University of Illinois at Urbana-Champaign in exchange for credit in an introductory Psychology course. All 15 gave informed consent based on the University of Illinois IRB protocol and reported normal or corrected-to-normal vision and color vision.

#### 4.1.2 Apparatus, Stimuli, and Procedure

We used the same apparatus and task as in previous experiments, but both the stimuli sizes and positions have changed. In Experiment 3, cues and targets appeared in one of eight positions (Figure 6) above and below the horizontal midline. There were four inner positions located 4° of visual angle from fixation and four outer positions located about 8.1° of visual angle from fixation. We scaled the stimuli to account for cortical magnification (Duncan & Boynton, 2003). The scaled inside stimuli subtended about 1.25° of visual angle and the scaled outside stimuli subtended about 2.2° of visual angle. Within a hemifield, the stimuli appeared at any of the four vertices of an imaginary square. The distance from any vertex to the within-hemifield vertices either directly horizontal or directly vertical is about 5.7° of visual angle. The inside upper vertex is 45° from the upper visual midline and the outside upper vertex is about 72° from the upper visual midline. The lower vertices are placed in symmetric locations. The timing parameters were the same as in previous experiments.

Experiment 3 again consisted of 50% valid and 50% invalid trials. The cues appeared equally often at all eight locations. When invalid, the target appeared equally often in one of the two locations within the same hemifield as the cue: either directly horizontal or directly vertical to the cue. So, 25% of trials were invalid-horizontal and 25% were invalid-vertical. Participants completed 448 total trials. (Note: slight differences in the number of trials between experiments are due to counterbalancing conditions with different numbers of levels.)

### 4.2 Results

#### 4.2.1 Inferential Statistics

Accuracy and response time data are presented in Table 2 and in Figure 7. The valid condition produced more accurate (Mean difference = 5.8%, SD = 6.0%, 95% CI of the mean = [2.3%, 9.2%], dz = 0.96, 95% CI of the effect size = [0.23, 1.57]) and faster responses (Mean difference = −60.7 ms, SD = 59.1 ms, 95% CI of the mean = [−94.86, −26.60], dz = −1.03, 95% CI of the effect size = [−1.38, −0.67]) than the invalid condition.

**Figure 7:**
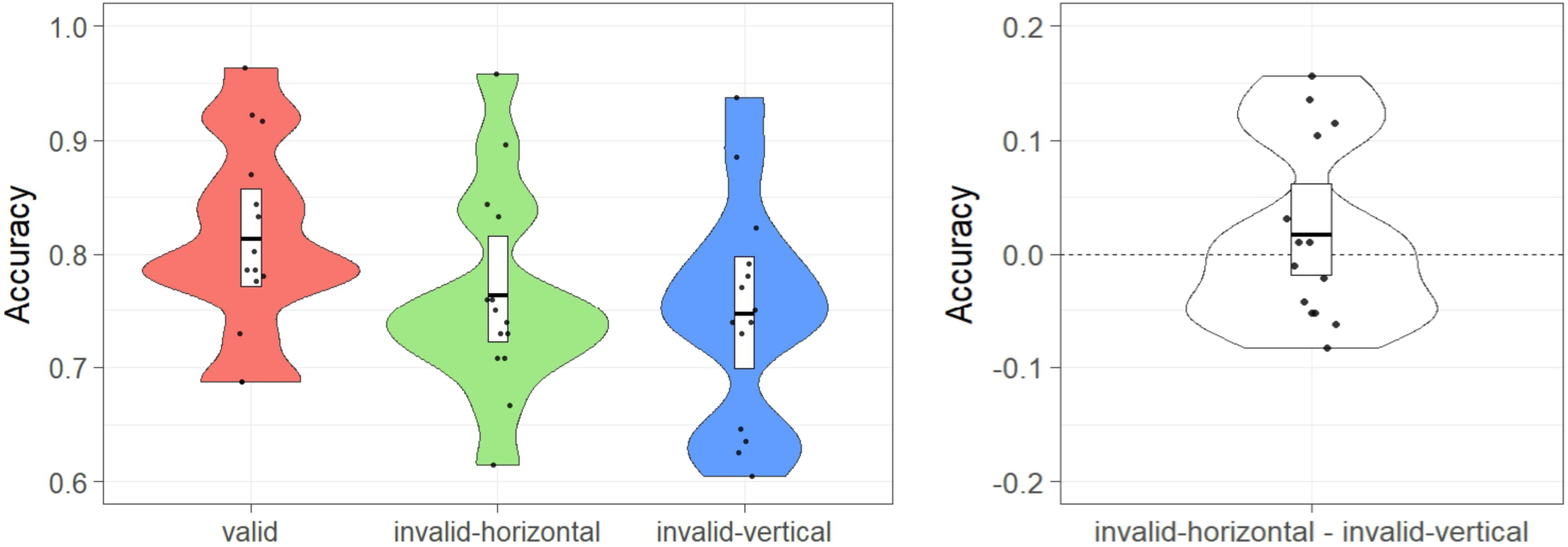
Accuracy from Experiment 3. Violin plot representation as in Figure 3. A) Accuracy as a function of condition. B) Participant-level accuracy differences between the invalid-different and the invalid-same conditions.

**Table 2.** Descriptive statistics for Experiment 3.

|  | Inner | Outer | Combined |
| --- | --- | --- | --- |
| Accuracy (%) |  |  |  |
| Valid | $78.5 \pm 1.0$ | $84.2 \pm 8.5$ | $81.3 \pm 9.7$ |
| Invalid-horizontal | $68.5 \pm 11.4$ | $84.3 \pm 7.8$ | $76.4 \pm 8.4$ |
| Invalid-vertical | $70.2 \pm 11.5$ | $79.2 \pm 11.5$ | $74.7 \pm 9.1$ |
| Response time (ms) |  |  |  |
| Valid | $744 \pm 126$ | $722 \pm 128$ | $733 \pm 125$ |
| Invalid-horizontal | $799 \pm 122$ | $769 \pm 123$ | $783 \pm 113$ |
| Invalid-vertical | $832 \pm 128$ | $775 \pm 77$ | $801 \pm 95$ |

Note: Mean ± Standard deviation

We again computed a difference score for each participant. We subtracted the participant mean in the invalid-vertical condition from the participant mean in the invalid-horizontal condition. This difference score will be positive to the extent that a participant was more accurate in the invalid-horizontal condition than in the invalid-vertical condition (Figure 7B). The group difference scores show a small but not statistically significant difference between the two conditions (M = 1.7%, SD = 8.0%, 95% CI of the mean = [−1.7%, 6.2%], dz = 0.22, bootstrapped 95% CI of the effect size = [−0.35, 0.76]). We again conducted a randomization test to determine whether the observed difference was greater than that expected due to chance. The procedure was the same as in previous experiments. Of the 100,000 iterations run, 41,310 produced a difference with an absolute value as great or greater than the observed difference (1.7%), resulting in p = 0.41. The same procedure was performed on the difference between valid and invalid conditions (collapsed across horizontal and vertical conditions) resulting in 380 difference scores with an absolute value as great or greater than observed, 5.8%, corresponding to p < .01

#### 4.2.2 Exploratory RT analyses

We analyzed the response time scores for both the invalid-horizontal condition and the invalid-vertical condition. We again computed difference scores. We subtracted each participant’s response time in the invalid-asymmetric condition from their response time in the invalid-vertical condition. The differences scores revealed a small benefit for the invalid-horizontal condition (M = 19 ms, SD = 47 ms). A randomization test (using the same procedure as above and uncorrected for multiple corrections/exploratory analyses) was conducted. Of the 100,000 iterations, 15392 produced a difference score with an absolute value as great or greater than the observed one (19 ms), p = 0.15. Thus, as in the accuracy analysis, although there was a numerical advantage for horizontally located targets over vertically located ones, this difference did not reach significance.

#### 4.2.3 Exploratory Accuracy analyses

The above results would suggest that the horizontal advantage does not occur within a hemifield. However, before concluding that the advantage reflects a horizontal hemifield advantage, there is another difference between Experiments 1 and 3 we must consider. In Experiment 1 the invalid location ultimately requires a shift away from fixation (although relative to the cued location attention would first have to move towards fixation). In Experiment 3, half of the invalid locations (the less eccentric ones) require only an inward shift of attention towards fixation, and the other half (the more eccentric locations) require an outward shift. If the horizontal advantage reflects something like a compensatory mechanism for having shifted to a peripheral location for the cue, then one might expect it to be more pronounced when the invalid location requires a shift towards a peripheral location.

Such a result was found by Barnas et al (Barnas & Greenberg, 2016) in one of their object-based attention experiments. Specifically, they found a statistically significant within-hemifield benefit for horizontal targets only when those targets appeared to the outside of the cues; there was a directionally similar but not statistically significant effect when target appeared to the inside of the cues. We investigated whether a similar pattern appeared in our data. Experiment 3 introduced a new condition where targets could appear to the inside of cues and where the direction of shifting attention was to the inside (toward fixation). We thus wanted to compare shifts of attention toward fixation to those away from fixation.

We also observed a large difference in performance between targets appearing in the four locations closer to fixation (4° of visual angle from fixation) compared to targets appearing in the four spots farther away from fixation (8.1° of visual angle). In the valid condition, accuracy for targets closer to fixation was lower (78.5%) and response times slower (744 ms) than for targets farther away from fixation (84.4% and 722 ms), suggesting that our cortical magnification manipulation overcompensated for eccentricity. This meant we could not cleanly compare horizontal shifts away from a cued location to vertical shifts away from the same location because each shift would land on a target at a different eccentricity. We thus conditioned not on cue location, but on target location (inner versus outer); that is, we compared performance at a given target location when the preceding cue was horizontal to the target to performance when the preceding cue was vertical to the target. Accuracy and RTs separated by whether the targets appeared in the inner positions or outer positions are presented in Figure 8 and Table 2. We again computed a difference score for each participant. We subtracted the participant mean in the invalid-vertical condition from the participant mean in the invalid-horizontal condition separately for the targets located closer to fixation (Shift-IN) and the targets located further from fixation (Shift-OUT). These difference scores will be positive to the extent that a participant was more accurate in the invalid-horizontal condition than in the invalid-vertical condition (Figure 8). The group difference scores for horizontal targets when they were located closer to fixation were not statistically significant from zero (M = −1.8%, SD = 12.8%, 95% CI of the mean = [−7.7%, 5.3%], dz = −0.14, bootstrapped 95% CI of the effect size = [−0.74, 0.46]). However, when the targets were located further for fixation (outward from the cue), the horizontal benefit just reached significance (M = 5.2%, SD = 9.2%, 95% CI of the mean = [0.009%, 10%], dz = 0.57, bootstrapped 95% CI of the effect size = [0.02, 1.1]). We note, however, that a direct comparison of the horizontal advantage for targets located closer to fixation (invalid-in) and the targets located further from fixation (invalid-out) did not reach significance (M = −0.7%, SD = 0.16%, 95% CI of the mean = [−0.160, 0.020], dz = −0.45).

**Figure 8:**
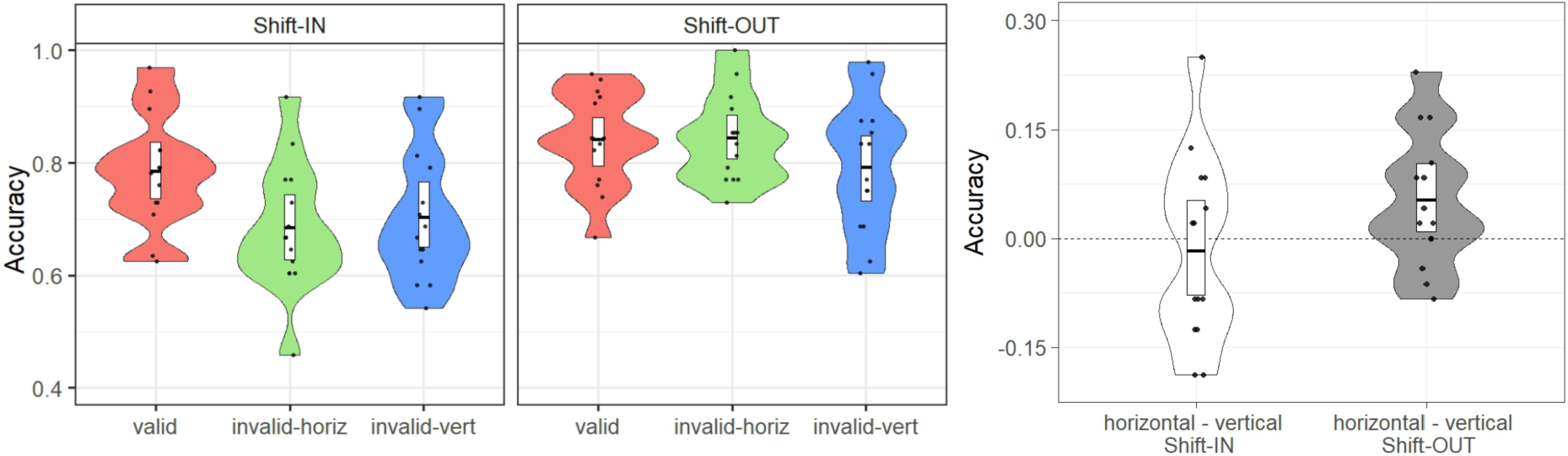
Accuracy from Experiment 3 as a function of shift direction. Violin plot representation as in Figure 3. A) Accuracy as a function of condition. B) Participant-level accuracy differences between the invalid-horizontal and the invalid-vertical conditions. Shift-IN denotes that the target was located at a position close to fixation while Shift-OUT denotes that a target was located at a position far from fixation.

Randomization tests (using the same procedure as above) showed similar results. For the cases in which the targets were close to fixation, 58,212 out of the 100,000 iterations produced a difference score with an absolute value as great or greater than the observed one (−1.8%), p = 0.58. For the cases in which the targets were far from fixation, 4,930 out of the 100,000 iterations produced a difference score with an absolute value as great or greater than the observed one (5.2%), p = 0.049.

### 4.3 Discussion

The results of Experiment 3 indicate that the horizontal advantage is greatly diminished when measured within a hemifield. Such a finding could reflect a horizontal but opposite hemifield advantage. By horizontal we mean that the target must still be located along a horizontal trajectory from the cue; Experiment 2 rules out a general effect of the opposite hemifield/hemisphere. However, exploratory analyses also raise the possibility that the horizontal advantage is obtained when attention must move away from fixation. A significant horizontal advantage was obtained only when the targets were located in the more peripheral locations, although the difference between peripheral and more foveal targets did not reach significance in this experiment. A similar asymmetry was found by Barnas et al. (2016) in an object-based attention paradigm. Such a pattern raises the possibility that the horizontal advantage reflects a compensatory mechanism of having focused attention on a location (as opposed to adopting a more global window of attention). When attention is focused locally, perhaps there is a compensatory tendency to also increase sensitivity to onsets in the periphery, locations that are already disadvantaged due to their eccentricity. This hypothesis, however, does not explain why the advantage is specific to horizontally located targets, since the vertical location is equally eccentric in our designs. This potential asymmetry will also need to be further evaluated with sufficient power to detect a difference between inner and outer targets. For now, a better synopsis of the results is that the evidence is weak that a horizontal advantage exists within a hemifield.

## 5. Experiment 4

Although we have ruled out the horizontal-vertical anisotropy and hemifield load as possible explanations, we still do not know why we observed a horizontal advantage. As mentioned in the introduction, one possibility is that this effect is related to the eye-movement system. Horizontal eye movements are more common than vertical ones (Foulsham et al., 2008; Gilchrist & Harvey, 2006; Tatler & Vincent, 2008; VanRenswoude et al., 2016), and this eye-movement bias may have been inherited by the attention system.

We suggested above that the differences we observed between inward and outward shifts of attention may reflect a compensatory mechanism that would increase resources assigned to more eccentric as opposed to less eccentric locations. A compensatory mechanism combined with an eye-movement explanation could also explain the opposite hemifield results of Experiment 1. When the eye moves horizontally this would render locations in the opposite visual field even more eccentric and thus particularly disadvantaged with respect to acuity. It is possible then that the visual system compensates for this disadvantage by increasing the probability that it will reorient back towards that disadvantaged location. Because vertical eye-movements are less common, and because the vertical extent of the visual field is smaller, no such compensatory mechanism may have developed. Although display times in our experiment preclude eye-movements, the fact that the attention system is so strongly linked to eye-movements may mean that the same compensatory mechanisms are at work even when the eye remains fixed and attention covertly moves.

The superior colliculus (SC), a subcortical structure involved in generating saccades and in exogenous orienting of attention, could be one candidate for enacting such a compensatory mechanism, or the horizontal bias more generally. Reflexive horizontal saccades have been shown to activate the SC as well as other brainstem nuclei in human (Linzenbold, Lindig, & Himmelbach, 2011). Rafal and colleagues (1988) assessed covert movements of attention in progressive supranuclear palsy (PSP), a degenerative disorder that effects subcortical nuclei including the SC. Patients were cued to one of four locations on the horizontal or vertical meridian. They found abnormalities in their response to the cues in both directions, when compared to the control subjects, but they emphasized a deficit in the vertical direction due to a reduced validity effect (invalid – valid) there (45ms) compared to in the horizontal direction (141ms). However, closer inspection of the components of the validity effect show that orienting to the cue (valid condition) was numerically faster in the horizontal direction (787ms) than the vertical (863ms), but that re-orienting to an invalid location was associated with a larger cost (i.e. validity effect) in the horizontal direction (141ms) than the vertical (45ms). In other words, the patients had particular trouble in reorienting attention in the horizontal direction, the condition in our experiment that shows superior reorienting relative to the vertical. These results raise the possibility that the SC may be involved in reorienting in the horizontal direction, and thus may be involved in the horizontal advantage we observed.

Luckily, owing to a known asymmetry in the nasal versus temporal hemifields of the SC, it is possible to assess the contribution of the SC to our task (Lewis, Maurer, & Blackburn, 1985; Wilson & Toyne, 1970). The retinotectal pathway, connecting the retina to the SC, has a stronger contralateral representation with a weaker ipsilateral contribution. The consequence of this inequity is that the temporal hemifield (nasal hemiretina) is a stronger driver of saccades and saccade-related effects (Kristjánsson, Vandenbroucke, & Driver, 2004; Lewis et al., 1985; Rafal, Smith, Krantz, Cohen, & Brennan, 1990; Walker, Mannan, Maurer, Pambakian, & Kennard, 2000) and attention (Rafal, Calabresi, Brennan, & Sciolto, 1989; Rafal, Henik, & Smith, 1991) than the nasal hemifield (temporal hemiretina). If the SC plays a central role in the horizontal advantage, we have observed then we would expect the advantage to be stronger in the temporal hemifield than the nasal hemifield. The hemifield stimulated by our stimuli can be controlled by occluding one eye. For example, if the left eye is occluded then stimuli presented in the left visual field will project to the temporal hemifield (strong connection to the SC) whereas stimuli in the right visual field will project to nasal hemifield (weaker connection to the SC). That relationship reverses if the right eye is occluded. In Experiment 5, we ask participants to perform our task with either the right or left eye occluded, alternately across the experiment.

### 5.1 Materials and Methods

#### 5.1.1 Participants

Because we are now looking for a difference between our original horizontal advantage and one that we conservatively estimated could be two thirds to one half the size (in the nasal hemifield), we chose a sample size of 30, to give us 80% to detect an effect size of dz=0.53. Thirty participants (mean age = 19.0 years old, and 23 females) from the University of Illinois took part in this experiment in exchange for class credit. All participants had normal or corrected-to-normal vision. All participants gave informed consent based on the University of Illinois IRB approved protocol.

#### 5.1.2 Apparatus, Stimuli, and Procedure

The apparatus, stimuli, and spatial arrangement of the stimuli were the same as in Exp.1, except that the distance between participants and the computer screen was 59 cm, and the resulting distance between target stimuli and fixation was 4.4° and the distance between two vertices of the imaginary square was 6.2°, with each vertex marking the center of a possible cue or target location.

In this experiment, we adopted a 3 (condition: valid, invalid-horizontal, invalid-vertical condition) × 2 (hemifield: Temporal and Nasal hemifield) within-subject design. Participants were asked to put on a pair of goggles that fully occluded one eye. Halfway through the experiment they switched the occluding side to the other. The order of eye occlusion (left or right) was counterbalanced across participants. Because eye-occlusion resulted in a drop in accuracy, target duration was increased to 141.6 ms and cue duration was increased to 100ms. The experiment contained 10 experiment blocks of 64 trials each (5 blocks for Temporal hemifield, and 5 blocks for Nasal hemifield). Participants did not wear the goggles during the practice block (64 trials). All other details of the experiment remained same as in Experiment 1.

### 5.2 Results

#### 5.2.1 Inferential Statistics

Means and standard deviation for accuracy and RTs are presented in Table 3 for both the temporal and nasal hemifield, as well as their combination. Figure 9 shows the accuracy data as a function of condition and hemifield, as well as accuracy difference scores between invalid-horizontal condition and invalid-vertical condition, but this time separately for the temporal and nasal hemifield. Both temporal and nasal hemifield showed a significant horizontal advantage, greater accuracy in invalid-horizontal condition than in invalid-vertical condition (Temporal: M = 4.7%, SD = 9.1%, 95% CI of the mean = [1.8%, 8.2%], dz = 0.51, 95% CI of the effect size = [0.16, 0.82]; Nasal: M = 5.8%, SD = 6.4%, 95% CI of the mean = [3.6%, 8.1%], dz = 0.90, 95% CI of the effect size = [0.57, 1.27]).

**Table 3.** Descriptive statistics for Experiment 4.

|  | Temporal | Nasal | Combined |
| --- | --- | --- | --- |
|  | Accuracy (%) |  |  |
| Valid | 85.0 ± 7.4 | 85.0 ± 7.0 | 85.0 ± 7.1 |
| Invalid-horizontal | 71.2 ± 10.6 | 72.7 ± 10.1 | 71.9 ± 10.3 |
| Invalid-vertical | 66.5 ± 12.3 | 67.0 ± 12.9 | 66.7 ± 12.5 |
|  | Response time (ms) |  |  |
| Valid | 641 ± 79 | 634 ± 79 | 639 ± 79 |
| Invalid-horizontal | 727 ± 108 | 728 ± 97 | 727 ± 102 |
| Invalid-vertical | 730 ± 102 | 726 ± 103 | 728 ± 102 |

Note: Mean ± Standard deviation

**Figure 9:**
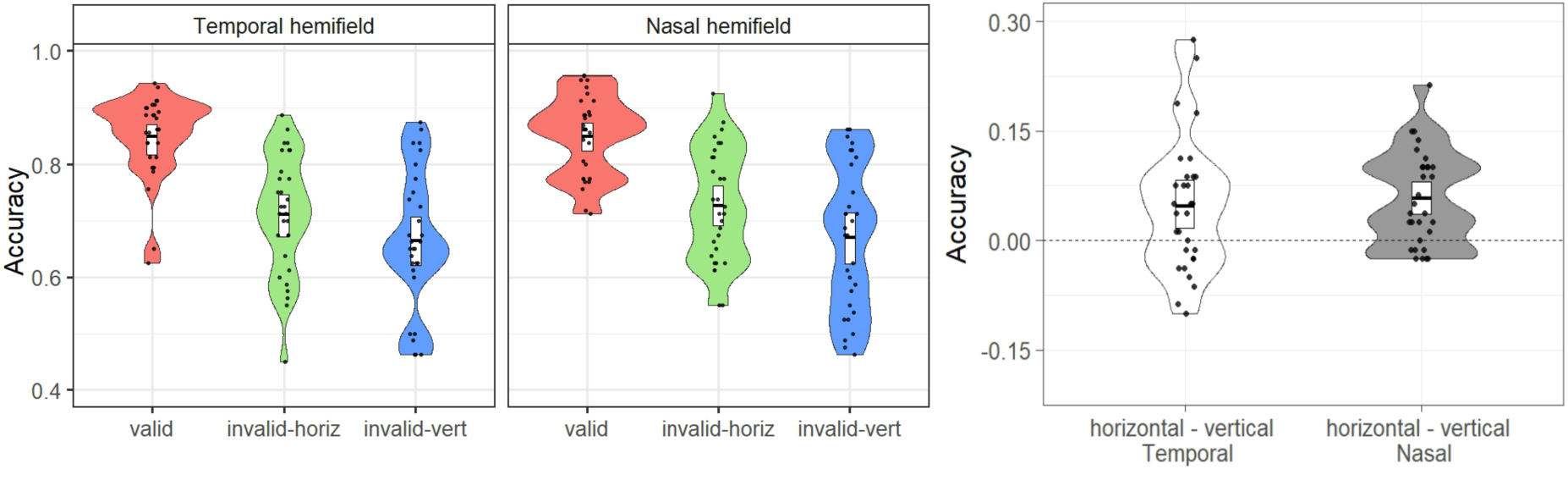
Accuracy from Experiment 4 as a function of hemifield. Violin plot representation as in Figure 3. A) Accuracy as a function of condition. B) Participant-level accuracy differences between the invalid-horizontal and the invalid-vertical conditions for temporal and nasal hemifield.

Randomization tests (using the same procedure as in Experiments 1-3) showed similar results. For the Temporal hemifield, of the 100,000 iterations, 789 produced a difference score with an absolute value as great or greater than the observed one, *p* < .01. For the Nasal hemifield, 4 of the 100,000 iterations produced a difference score with an absolute value as great or greater than the observed difference score, *p* < .001. Because the temporal and nasal hemifields produced virtually identical result, no follow-up inferential tests were made to assess hemifield differences. There was clearly no effect of hemifield. The same procedure was performed on the difference between valid and invalid conditions (collapsed across horizontal and vertical conditions) resulting in 0 difference scores with an absolute value as great or greater than that observed (i.e. 15.7%), corresponding to p < .001.

#### 5.2.2 Exploratory RT analyses

We computed the same difference scores for RTs as we did for accuracy, again separately for the temporal and nasal hemifield. No significant difference between invalid-horizontal and invalid-vertical condition was observed for either the temporal and nasal hemifield (Temporal: M = 3 ms, SD = 32 ms, 95% CI of the mean = [−8.3, 14.1], dz = 0.09, 95% CI of the effect size = [−0.28, 0.47]; Nasal: M = −2 ms, SD = 53 ms, 95% CI of the mean = [−20, 18], dz = −0.04, 95% CI of the effect size = [−0.42, 0.33]). Randomization tests showed similar results, with no difference between invalid-horizontal and invalid-vertical condition (Temporal: 61318/100000 was as great or greater than the observed difference, *p* = 0.622, Nasal: 81192/100000 was as great or greater than the observed difference, *p* = 0.817).

### 5.3 Discussion

We found no evidence of a differential horizontal advantage for the temporal and nasal hemifield. However, we did replicate the horizontal advantage, for both hemifields, using a much larger sample size; assuming an effect size of dz=1, we had 99.9% power to detect the difference in Experiment 4. The lack of a temporal/nasal asymmetry argues against a strong role for the SC in the observed horizontal advantage. However, these data do not rule out an eye-movement explanation of our effect. SC is only one component of the subcortical eye-movement system. For instance, studies show that central mesencephalic reticular formation (cMRF), which receives inputs from the SC, may be a more critical driver of horizontal saccades (Wang, Perkins, Zhou, Warren, & May, 2017). The cMRF then projects to the paramedian pontine reticular formation (PPRF). It is unclear how the SC asymmetry propagates through the system, especially given that the SC is modulated by the cortical eye-movement system (Fries, 1984; Segraves & Goldberg, 1987; Sommer & Wurtz, 2000; Wilson & Toyne, 1970) which does not show a similar nasal-temporal asymmetry (Sylvester, Josephs, Driver, & Rees, 2007).

It is thus still unclear why this horizontal advantage exists. Experiment 4 was motivated by the idea that the advantage may actually be an advantage in shifting attention, a hypothesis based on the fact that the oculomotor system and attention system are so intimately linked. It is also possible, however, that the horizontal advantage does not stem from shifts of attention per se, but from a pooling of attention in the opposite hemisphere (Tse et al., 2003). Transcallosal connections between visual hemispheres do not support such a view, as they appear to only connect visual hemifields in a narrow band around the vertical midline (Clarke & Miklossy, 1990; Rochefort, Buzás, Kisvárday, Eysel, & Milleret, 2007). Of course, it is possible that such an effect does not occur via direct transcallosal connections, but rather depends on feedback from higher visual areas (Ban, Yamamoto, Fukunaga, Nakagoshi, Umeda, Tanaka, & Ejima, 2006). Using fMRI, Ban and colleagues (2006) observed contextual activations from the opposite hemisphere of early visual cortex (V1-V3) but they were not symmetric across the vertical midline; instead, a diagonal symmetry (e.g. lower left visual field co-activated upper right visual field) was seen. Such a pattern of activity would predict strong between hemifield effects in our Experiment 2, which placed stimuli in the diagonal hemifields. However, no differences were found between the different hemifield condition and the same hemifield condition in that experiment. The only time a robust horizontal advantage was observed was when the target appeared in a symmetrical location in the opposite hemifield (Experiments 1 and 4).

Another possible argument against a shifting explanation of our effect is that our effects were confined to accuracy. Although there was a weak horizontal advantage in RTs for Experiment 1, that difference was clearly not significant with a higher-powered study (Experiment 4). We note that our study was designed to assess the effects of invalid target locations on accuracy, but if the underlying mechanism for our horizontal advantage in accuracy was a tendency to shift attention horizontally we would expect a parallel effect in RT. Instead, it seems that our results stem more from an enhanced representation of the target in a horizontal location from the cue, perhaps due to better selection of the target among flankers (Beck & Kastner, 2009; Desimone & Duncan, 1995). Of course, it is also possible that other factors influencing RTs (Duncan, 1980; Prinzmetal, McCool, & Park, 2005) are obscuring an RT difference, in which case it is still possible our advantage is related to a relative ease in moving attention horizontally compared to vertically.

Related to the pooling at symmetrical locations idea, it is also possible that attention is more easily elongated (to encompass both validly and invalidly cued locations) in the horizontal direction than the vertical direction. Because the cue is only valid 50% of the time it is possible that participants attempt to spread attention beyond the cued location and do so better in the horizontal direction. Some of the same justifications for a horizontal eye-movement bias might also justify a horizontal elongation of attention: the distribution of objects in the world tends to occur along the horizon and we may thus have learnt to distribute our attention in a similar fashion; our visual field, by virtue of having two horizontally displaced eyes, has a greater extent in the horizontal direction than it does in the vertical and attention maybe distributed to accommodate this fact. Although there are a number of just-so stories we could tell about a horizontal advantage, we know of no independent data, beyond the effect investigate here, that would corroborate the hypothesis that attention tends to be elongated more so in the horizontal direction than the vertical (when it is off the visual meridians, as it is here).

Finally, we should note that it is unclear whether a horizontal advantage will be observed for all stimuli and all cueing tasks. If we limit the cases to just those studies that demonstrated a horizontal advantage that was not confounded with the HVA, then it has been seen for exogenous cueing in both a Posner cueing task (here) and an object-based but exogenous cueing task (Barnas & Greenberg, 2016; Pilz et al., 2012) and for discriminating upright from inverted faces (here), discriminating at T from an L (Pilz et al., 2012), and detecting a T or an L (Barnas & Greenberg, 2016; Pilz et al., 2012). Zénon et al., (2009) also reported a horizontal advantage that was not confined to the horizontal meridian, using a visual search and letter detection task, but their target placements were such that they were still within 26° of the horizontal meridian, a location that still shows some benefit compare to stimuli locate on or near the vertical meridian. Further work is needed to understand the boundary conditions of this advantage.

Although the source of the horizontal advantage that we observed is unknown, our results still have important implications for a number of attention paradigms. If recruiting attention via an exogenous cue leads to enhanced processing in the hemifield opposite to the cue, this would mean that we might be underestimating the effect of a cue in a ubiquitous attention paradigm, the Posner cueing task (Posner, 1980). The Posner cueing task is one of the canonical demonstrations that covert attentional shifts can speed up processing of stimuli at validly cued locations and slow down processing of stimuli at invalidly cued locations (Posner, 1980). Typically, the task involves participants fixating in the center of a screen with the cues and targets appearing in symmetrical locations in the opposite hemifield, sometimes along the horizontal meridian and sometimes in an imaginary square around fixation, as in our design. The speeding of responses in the valid condition is taken to reflect the covert deployment of attention to the cued location and the difference between the valid and invalid conditions is taken to represent the cost having to re-deploy attention from the cued location to the uncued location. If, however, a location to the left of fixation is cued and there is a subsequent pooling of attention on the right side of the screen, responses to the right side (in this case the invalid condition) might not reflect the true cost of re-deploying attention to the right side because attention might already partly be there.

Similarly, our results draw into question an object-based attentional cueing effect. In Experiments 1 and 4 we observed a large benefit for horizontal targets using a display type like those in the classic object-based attention tasks (Egly et al, 1994). Interestingly, Pilz et al (2012) used a similar version of the classic task and found evidence for a same-object benefit only when same-object benefit was confounded with the horizontal advantage. In fact, they found evidence for a between-object benefit (the opposite of the classic effect) when the between-object benefit was confounded with the horizontal advantage. In other words, it seems like the object-based effect in Pilz et al (2012) might be partially or completely explained by the horizontal advantage we observed here. Our results correspondingly indicate that the horizontal advantage might be a general feature of spatial attention, rather than a property of object-based attention.

## 6. Conclusions

We observed a horizontal advantage for targets placed in a symmetric location in the opposite hemifield/hemisphere to the cue. This advantage did not depend on the HVA since the stimulus locations were chosen to appear outside the zone of the asymmetry. The horizontal advantage was not obtained when the stimulus locations were rotated 30° relative to those in Experiments 1 and 4, such that the stimuli were no longer horizontally symmetric. Nor did the effect occur when the targets were place in a horizontal location within the same hemifield as the cue. However, there was some evidence that there might be a horizontal advantage for movements away from fixation but within a hemifield, that is, when the target was presented more peripherally than the cue within the same hemisphere. These results will need further replication and examination to fully assess the boundary conditions for the horizontal advantage. Although neither the cause of this effect nor its boundaries are yet clear, our data raise the important possibility that we very often underestimate the effects of attentional cueing due to a pervasive bias across many tasks to place targets in symmetrical locations across the vertical midline.

## Acknowledgements

These experiments were included in a dissertation submitted by John Clevenger in partial fulfillment of the requirements for a doctoral degree at the University of Illinois. This research did not receive any specific grant from funding agencies in the public, commercial, or not-for-profit sectors.

## Footnotes

1 Although Pilz et al.’s (2012) data is consistent with a horizontal advantage and no object-based effect, we are not arguing that there are no object-based effects in general. Indeed, Barnas & Greenberg (2016) have evidence of both an object-based effect and a horizontal advantage.

